# Verifying molecular clusters by 2-color localization microscopy and significance testing

**DOI:** 10.1101/847012

**Authors:** Andreas M. Arnold, Magdalena C. Schneider, Christoph Hüsson, Robert Sablatnig, Mario Brameshuber, Florian Baumgart, Gerhard J. Schütz

## Abstract

While single-molecule localization microscopy (SMLM) offers the invaluable prospect to visualize cellular structures below the diffraction limit of light microscopy, its potential could not be fully capitalized due to its inherent susceptibility to blinking artifacts. Particularly, overcounting of single molecule localizations has impeded a reliable and sensitive detection of biomolecular nanoclusters. Here we introduce a 2-Color Localization microscopy And Significance Testing Approach (2-CLASTA), providing a parameter-free statistical framework for the analysis of SMLM data via significance testing methods. 2-CLASTA yields p-values for the null hypothesis of random biomolecular distributions, independent of the blinking behavior of the chosen fluorescent labels. We validated the method both by computer simulations as well as experimentally, using protein concatemers as a mimicry of biomolecular clustering. As the new approach it is not affected by overcounting artifacts, it is able to detect biomolecular clustering of various shapes at high sensitivity down to a level of dimers.

## Introduction

Single Molecule Localization Microscopy (SMLM) has boosted our insights into cellular structures below the diffraction limit of light microscopy^1^. Common to all SMLM variants is the stochastic switching of single dye molecules between a bright and a dark state. Conditions are chosen such that only a marginal portion of the molecules is in the bright state, so that single molecule signals are well separated on each frame. The final superresolution image is reconstructed from the localizations of all single molecule signals.

Researchers have been particularly intrigued by the possibility to determine the spatial distribution of biomolecules in their natural environment, in most cases the intact cell. For example, models for cellular signaling are crucially affected by the spatial organization of receptor and downstream signaling molecules at the plasma membrane^2,3^. Indeed, application of SMLM to various plasma membrane proteins revealed the presence of nanoclusters to different degrees^4^. More recently, however, concerns were raised that the stochastic activation process of the fluorophores, along with the presence of more than one dye molecule per labeled biomolecule, may lead to multiple observations of the same biomolecule in the superresolution image^5,6^. Different attempts were undertaken to approach this problem^5,7–11^, e.g. by merging localization bursts into one localization^12^, by analyzing the number of blinking events per localization cluster^10,11^, or by evaluating the spatial spread of the localization clusters^7^. A disadvantage of existing methods is the requirement of user-defined parameters^7,12^ or additional experiments to characterize the blinking statistics of the chosen fluorophores^10,11^. Recently, we came up with a parameter-free method to identify global protein clustering based on a label titration approach^8^ (see also^9^), however, in case of faint bimolecular clustering the discrimination is difficult and rather subjective. Taken together, it would be helpful to provide a parameter-free quantitative assessment for the reliability of the statement, whether biomolecular nanoclusters occur in an image or not.

Here we present a method to assess biomolecular nanoclustering in SMLM via p-values in the framework of statistical significance tests, termed 2-Color Localization microscopy And Significance Testing Approach (2-CLASTA). The idea is to target the same biomolecule of interest with different fluorescent labels, determine the localizations in the respective color channels, and calculate the nearest neighbor distances between them. The test compares the nearest neighbor distances for the recorded data with the distances from a random distribution of biomolecules calculated from the measured data. As an output, the method provides a p-value for the null hypothesis that the experimental data set corresponds to an underlying biomolecular distribution, which is not significantly different from a completely random distribution as described by a spatial Poisson process. In this respect, 2-CLASTA differs from existing approaches, which typically aim at determining quantitative parameters before actually testing the mere presence of biomolecular clusters. The method is parameter-free and does not require any additional measurements. We validated the method experimentally in cells expressing artificially clustered proteins by showing that sizes down to 2 molecules per cluster can be reliably detected.

## Results

### Testing the null hypothesis of a random biomolecular distribution

In principle, labeling the biomolecule of interest in two different colors yields different two-color SMLM images for a random versus a clustered biomolecular distribution (**Fig. 1a**). Both images show clear clustering of localizations in each of the color channels due to multiple observations of single dye molecules. The localization clusters of different color, however, correlate only in case of an underlying clustered distribution of biomolecules. As a quantitative measure of this correlation we used the empirical cumulative distribution function, *cdf*, of the nearest neighbor distance, *r*, between the localizations of the two different color channels. Importantly though, *cdf*(*r*) not only depends on the spatial distribution of the labeled biomolecule. Particularly, the blinking statistics of the fluorophore and the number of dye molecules conjugated to the biomolecule of interest affect the distribution functions. Since these parameters are commonly unknown, the different contributions to *cdf*(*r*) are difficult to disentangle.

**Figure 1.**
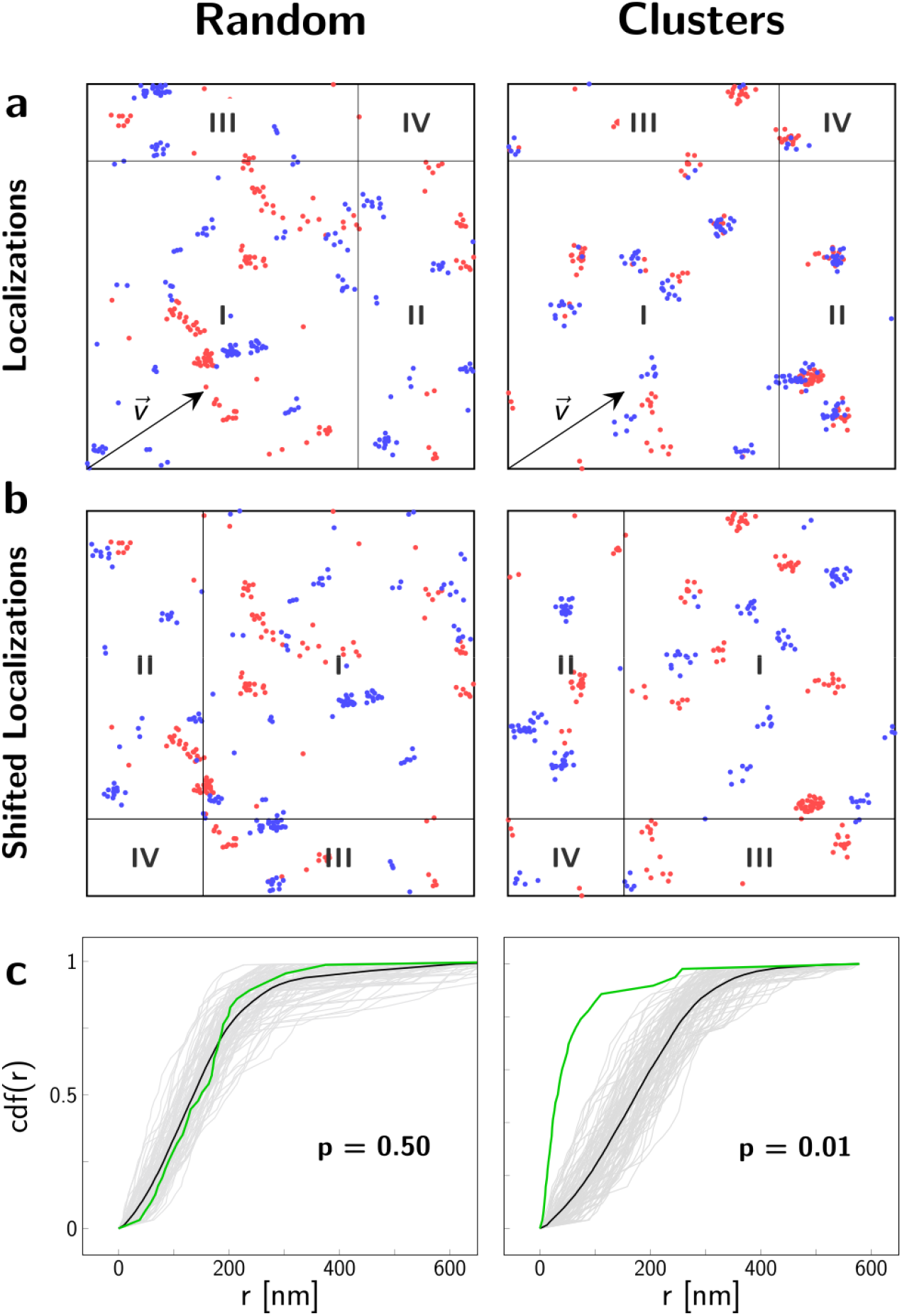
Analysis of localization maps with 2-CLASTA. (**a**) Simulated two-color localization maps for a random (left column) and a clustered (right column) distribution of biomolecules. Images show a 2 x 2 μm^2^ region. For the simulation of blinking we used experimental data obtained for SNAP_488_ (blue channel) and SNAP_647_ (red channel). (**b**) Shifting all localizations of the blue color channel by the shift vector 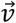 breaks correlations between the two color channels. (**c**) The cumulative distribution function of nearest neighbor distances, r, between the two color channels is plotted in green for the localization data shown in (a). *cdf_rand_* of N=99 control curves, generated with randomly chosen toroidal shifts, are depicted in light gray. The mean of all control curves is shown in black. From the rank of the curves, we calculated a p-value p=0.50 for the random case, and p=0.01 for the clustered case.

To analyze the data, we hence opted for a strategy which is independent of prior information on label properties. The idea is to determine a randomized distribution function *cdf_rand_*(*r*) for a scenario in which correlations between the two color channels are broken, by directly using the experimental data contained in the original SMLM recording. Our approach is similar to a goodness-of-fit test, in which the experimental data are compared with Monte Carlo-simulated control data sets using a global deviation measure for calculation of a p-value^13^.

In order to construct a randomized two color data set we transformed the localizations of one color channel and calculated their nearest neighbor distances to the untransformed localizations of the other color channel. For the transformation we used a toroidal shift, which breaks potential correlations between the two color channels^14^ (**Fig. 1b**). The resulting *cdf_rand_*(*r*) implicitly accounts for the correct blinking statistics and degree of labeling, and can hence be taken as ground truth for the situation of two uncorrelated images, irrespective of their univariate clustering that may be present in each color channel itself. Ideally, for a completely random protein distribution the cumulative density functions are equal (*cdf*(*r*) = *cdf_rand_*(*r*)), whereas for a non-random distribution they are not (*cdf*(*r*) ≠ *cdf_rand_*(*r*)). Note that *cdf_rand_*(*r*) does not need to correspond to a truly random distribution of molecules.

For the statistical assessment, we compared the original empirical *cdf*(*r*) with a set of N=99 realizations of *cdf_rand,i_* (*i* = 1,…, *N*) for random choices of the toroidal shift vector 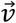 (**Fig. 1c**), and ranked the summary statistics *g_data_* of the original curve (green) with respect to the control curves (gray) (see Methods). Since we are interested in nanoclustering of biomolecules, we determined the one-sided p-value by ranking the original *cdf*(*r*) with respect to all calculated *cdf_rand,i_*; the rank is measured in descending order (note that the method also allows for assessing biomolecular repulsion by calculating the rank in ascending order). In practice, prior knowledge on cluster sizes can be taken into account e.g. by constraining the analysis to short distances. For this, we introduced a parameter *r_max_*, which should be chosen close to the minimum of the localization errors and the expected cluster size. Here we ignored prior knowledge and set *r_max_* →∞, if not mentioned otherwise.

Naturally, the p-value as defined here is limited to discrete numbers with steps of 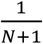, which also defines the minimum p-value obtainable with this method. As expected, the p-value is uniformly distributed in the interval [0,1], when testing realizations of the null hypothesis against the null hypothesis itself (**Fig. S2**). Hence, this p-value allows for the correct interpretation of the significance level α as the probability of falsely rejecting the null hypothesis. α can also be interpreted as the inevitable false positive rate for the erroneous detection of overcounting-induced clustering for a random distribution of biomolecules. Taken together, by offering an appropriate significance test, 2-CLASTA is hardly susceptible to the inadvertent interpretation of localization clusters as biomolecular nanoclusters.

On the other hand, it is crucial that the test is sufficiently sensitive to detect even faint spatial biomolecular clustering. We assessed the sensitivity (also frequently termed power) of 2-CLASTA for two clustering scenarios: i) biomolecular oligomerization (dimers, trimers, and tetramers), and ii) spatially extended clusters with varying load. The spatial distribution of the biomolecules and the according localization maps were generated with Monte Carlo simulations and evaluated with 2-CLASTA. We quantified the test performance via the sensitivity defined as 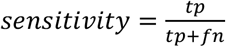, with *tp* denoting the true positives (here defined as correctly detected clustering) and *fn* the false negatives (here defined as erroneously missed clustering). We used a significance level α=0.05 in the following.

### Sensitivity to detect biomolecular oligomerization

We first assessed the sensitivity of 2-CLASTA to detected different degrees of oligomerization. For this, we simulated 10 x 10 μm^2^ sized images containing randomly distributed dimers, trimers, or tetramers, assigned labels of the two colors with the according blinking statistics, and added localization errors. Each image can be considered as a realization of a two-color superresolution experiment. The images were analyzed by 2-CLASTA, yielding a p-value for each image and the sensitivity for each parameter set. We showcase the performance of the method with an “ideal” scenario, which lacks the presence of unspecific signals, and assumes a degree of labeling of 100%. In a real-life experiment, however, unspecifically bound fluorophores and background signals may be present in the final localization maps, which may affect the obtained statistics. We hence also analyzed a more “realistic” scenario, for which we added 5 unspecifically bound dyes per μm^2^ in each color channel, and 1 or 2 unspecific background signals in the red or blue color channel, respectively; the characteristics of the unspecific background signals were experimentally determined on unstained cells. For the “realistic” case, we further assumed a reduced degree of labeling of 40%. If not specified otherwise, the degree of labeling for both colors was simulated to be balanced.

We were first interested in the total number of biomolecules per image that are required for a reliable detection of oligomerization. Already low numbers of biomolecules of ~1,000 per image (corresponding to 10 molecules per μm^2^) allow for a sensitive detection even of dimerization, both for the “ideal” and the “realistic” scenario (**Fig. 2a**). As expected, the sensitivity is somewhat reduced with decreasing degree of oligomerization: this is a consequence of the reduced fraction of oligomers carrying two different labels, particularly for the “realistic” scenario. For the following simulations, we used 7,500 molecules per image (75 molecules/μm^2^). We next tested the influence of a reduced labeling efficiency. In general, sensitivity was found to be high even down to a labeling degree of ~20% (**Fig. 2b**). To test the influence of different blinking statistics or of multiple dye molecules per label, we used experimentally derived blinking statistics for SNAP and SNAP (**Fig. S1**) as well as the blinking behavior of PS-CFP2 (blue channel) and an Alexa Fluor 647-conjugated antibody (KT3_647_, red channel)^15^ for our simulations, yielding virtually identical results (**Fig. S3**).

**Figure 2.**
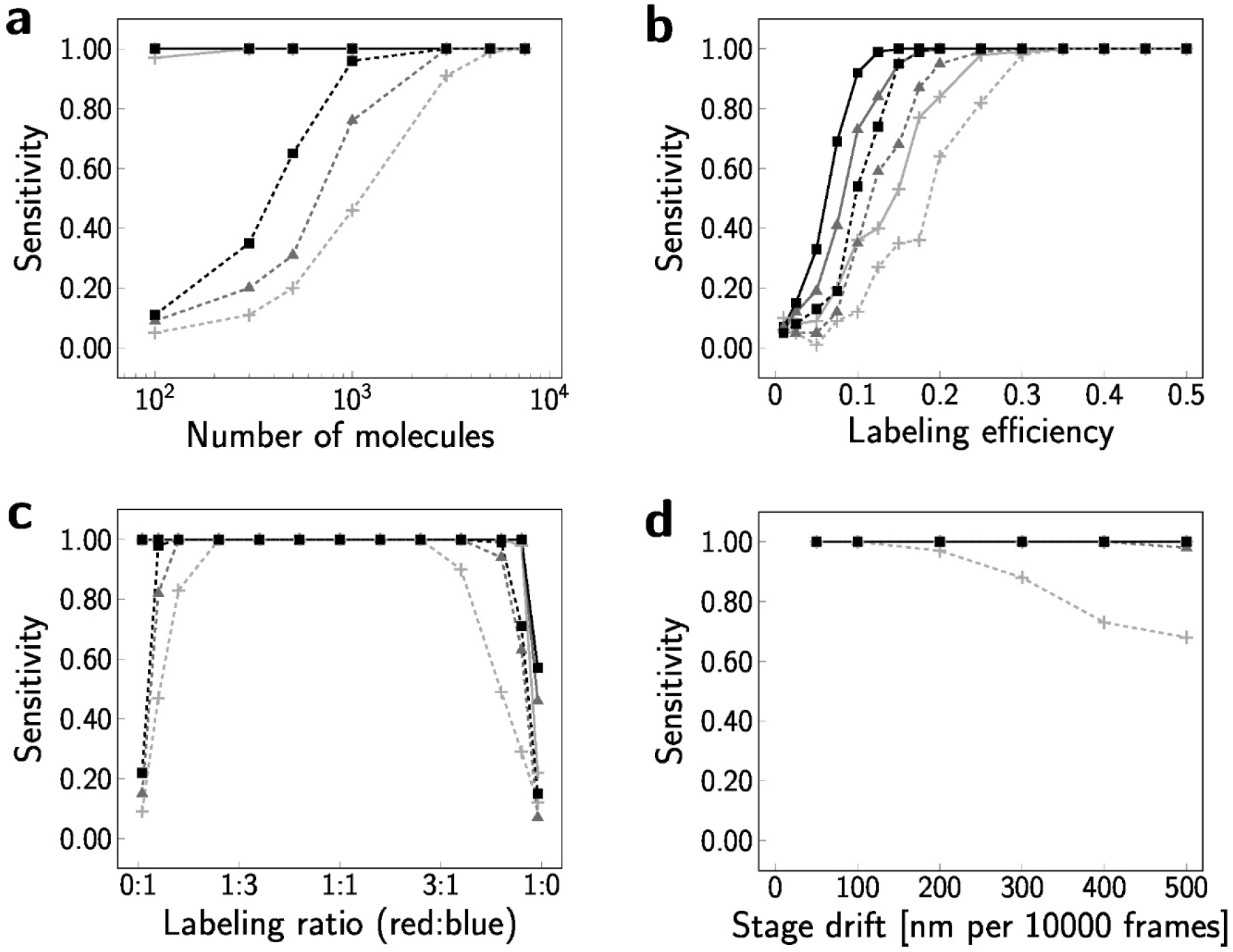
Robustness of 2-CLASTA for the detection of different degrees of oligomerization. To assess the influence of individual parameters we determined the sensitivity as a function of the number of molecules (**a**), the labeling efficiency (**b**), the labeling ratio (**c**), and directional stage drift (d). We simulated dimers (✚), trimers (▲) and tetramers (■), both for the “ideal” (solid line) and the “realistic” scenario (dashed line). For panel (**d**), virtually all simulated scenarios yielded a sensitivity of 1. If not varied in the respective subpanel, parameters in all simulations were set to a molecular density of 75 molecules/μm^2^, a labeling efficiency of 40% for the real case and 100% for the ideal case, a labeling ratio of 1:1, and no stage drift.

While the fraction of each label should ideally be kept around 50%, we found the sensitivity to remain high also for unbalanced labeling (**Fig. 2c**). Next, we also tested the influence of randomly distributed unspecific labels added onto the simulated oligomer distributions, yielding only marginal influences (**Fig. S4a**). Further, the magnitude of the localization errors hardly affects the obtained results (**Fig. S4b**). Interestingly, an assessment of the influence of stage drift showed that drift-velocities of up to 500 nm over 10.000 frames hardly affected the test sensitivity (**Fig. 2d**). This is not unexpected, as drift hardly diminishes the correlations between the two color channels in an experiment performed at alternating laser excitation. Finally, we evaluated the influence of different values of *r_max_* on the sensitivity of the method, yielding only minor effects (**Fig. S5**).

### Sensitivity to detect areas of enrichment or depletion of biomolecules

As a second realization of a non-random spatial distribution of biomolecules we considered spatially extended circular domains, the centers of which were randomly distributed across a two-dimensional plane. Molecules were placed either inside or outside of the domains, which thereby represent areas enriched or depleted in biomolecules compared to the surface density outside of the domains. To facilitate comparison with our previously published approach^8,15^, we used here the same parameter settings for assessing the performance of 2-CLASTA: we varied the domain radius between 20nm and 150nm, the domain density between 3 and 20 domains per μm^2^, and the fraction of molecules in domains between 20% and 100% (**Fig. S6**). The overall density of biomolecules was kept constant at 75 molecules per μm^2^.

In general, virtually all scenarios with a substantial heterogeneity in the lateral distribution of the biomolecule can be detected by 2-CLASTA (**Fig. 3a** and **Fig. S7**): both biomolecular clustering (top right corner) and exclusion areas (bottom left corner) yield a high level of correctly identified scenarios. In particular, the new method even outperforms our previous approach based on label titration, as can be seen by comparing the new figures with the respective plots from our previous paper (Supplementary Figure 5 and 6)^15^.

**Figure 3.**
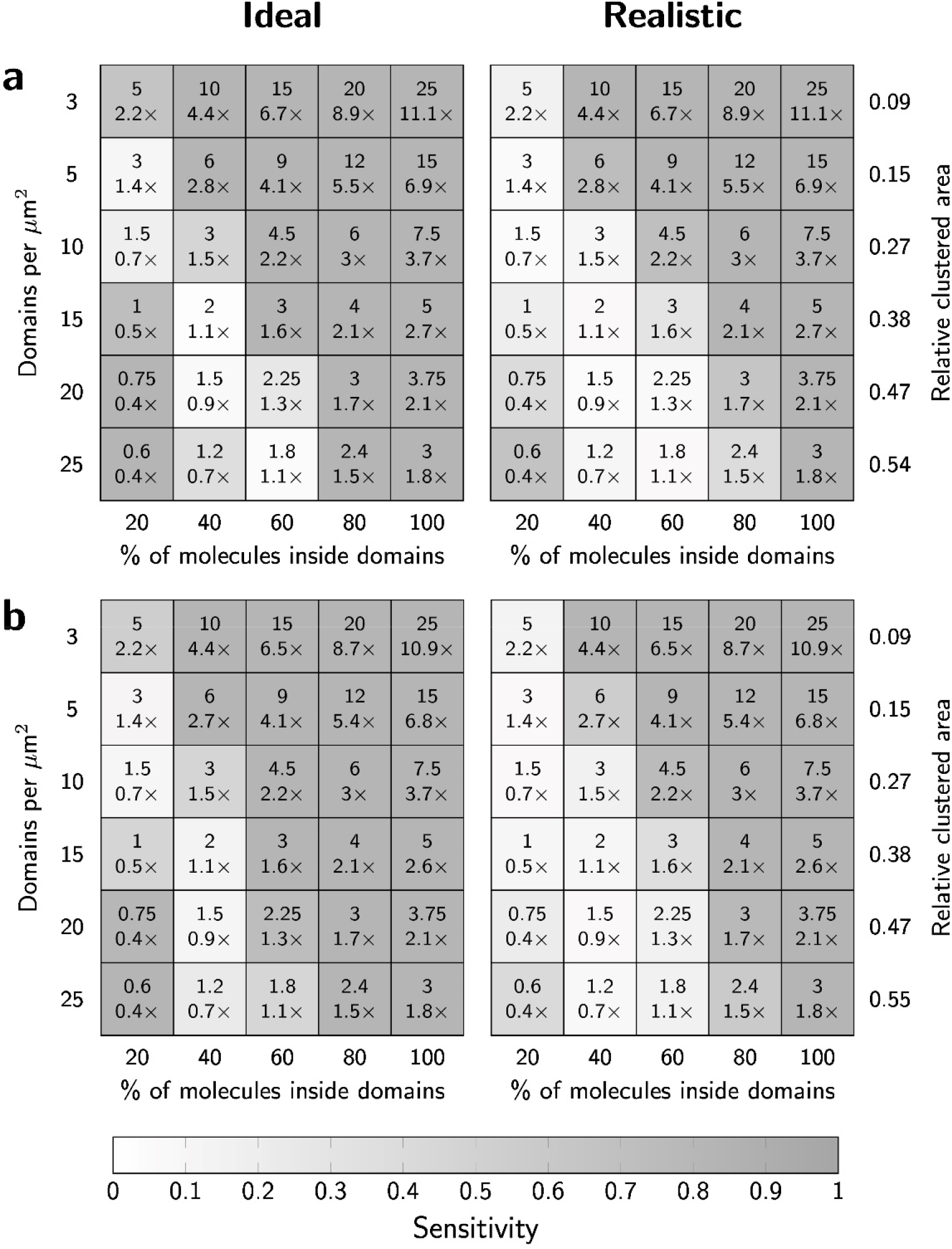
Sensitivity of 2-CLASTA to detect protein enrichment or depletion. (**a**) We determined the sensitivity of 2-CLASTA for varying densities of circular domains and percentage of molecules inside the domains. Data are shown for a cluster radius of 100 nm for the “ideal” case and the “realistic” case (see **Fig. S7 & S8** for other cluster radii). (**b**) The sensitivity for the detection of rectangular clusters with a size of 80 x 400 nm2 is shown for the ideal case and the “realistic” case. Numbers in individual fields indicate the average number of molecules per domain, and the relative enrichment or depletion of molecules compared to a random distribution with identical average density. The gray sale indicates the fraction of scenarios with a p-value below the significance level α=0.05, reflecting the sensitivity. Each field corresponds to 100 independent simulations.

The diagonal in **Fig. 3a** represents scenarios, in which the biomolecular concentration inside the domains is similar to the concentration outside of the domains. In other words, these situations correspond to random distributions of biomolecules, which – if detected – would lead to false positive results. Per definition, a random distribution leads to a false positive rate that is identical to the chosen level of significance (here α=0.05). Indeed, for scenarios corresponding to identical biomolecular densities (±10%) inside versus outside the domains we obtained sensitivity values close to α, hence reaching the principal limit for analyzing a statistical data set.

Also here, we simulated a more “realistic” scenario as defined above, yielding similar results as for the ideal scenarios (**Fig. 3a** and **Fig. S8**). To assess whether the use of different fluorescent labels with altered photophysical properties affect the results, we repeated the simulations both for the “ideal” and the "realistic” case using the blinking statistics derived previously for a multi-labelled antibody and the photoactivatable protein PS-CFP2^15^, yielding virtually unchanged results (**Fig. S9**). Finally, we tested the algorithm on rectangular clusters of 80 x 400 nm^2^ size (**Fig. 3b**), yielding similar sensitivity as for circular domains of the same area coverage. In conclusion, the new approach allows for reliable detection of even faint biomolecular clustering, and is not susceptible to false positives due to overcounting artifacts.

### Experimental validation

For experimental validation of the 2-CLASTA approach, we mimicked protein monomers and oligomers by concatemers of SNAP-tags with 1 to 4 subunits. These concatemers were anchored in the plasma membrane of HeLa cells via a glycosyl-phosphatidylinositol-(GPI-) anchor. For example, SNAP-concatemers of 4 SNAP-tag subunits would correspond to tetrameric protein oligomers. Clustering of the GPI anchor *per se* is not expected^7,8^. All experiments were performed at similar labeling densities of SNAP-Surface Alexa Fluor 488 (SNAP_488_) and SNAP-Surface Alexa Fluor 647 (SNAP_647_). dSTORM experiments were performed at alternating excitation, yielding superresolution images of the two color channels (**Fig. 4a**). For each concatemer, we recorded ≥25 cells, and determined the according p-value for the null-hypothesis of a random protein distribution, as described above (**Fig. 4b**). For SNAP-monomers, we observed a uniform distribution of p-values in the interval [0,1], hence providing no indication for a non-random distribution. In contrast, dimeric, trimeric, and tetrameric SNAP-constructs yielded clear deviations from a uniform distribution, with a substantial peak at low p-values. This reflects the expected signature for an underlying non-random distribution of SNAP-tags.

**Figure 4.**
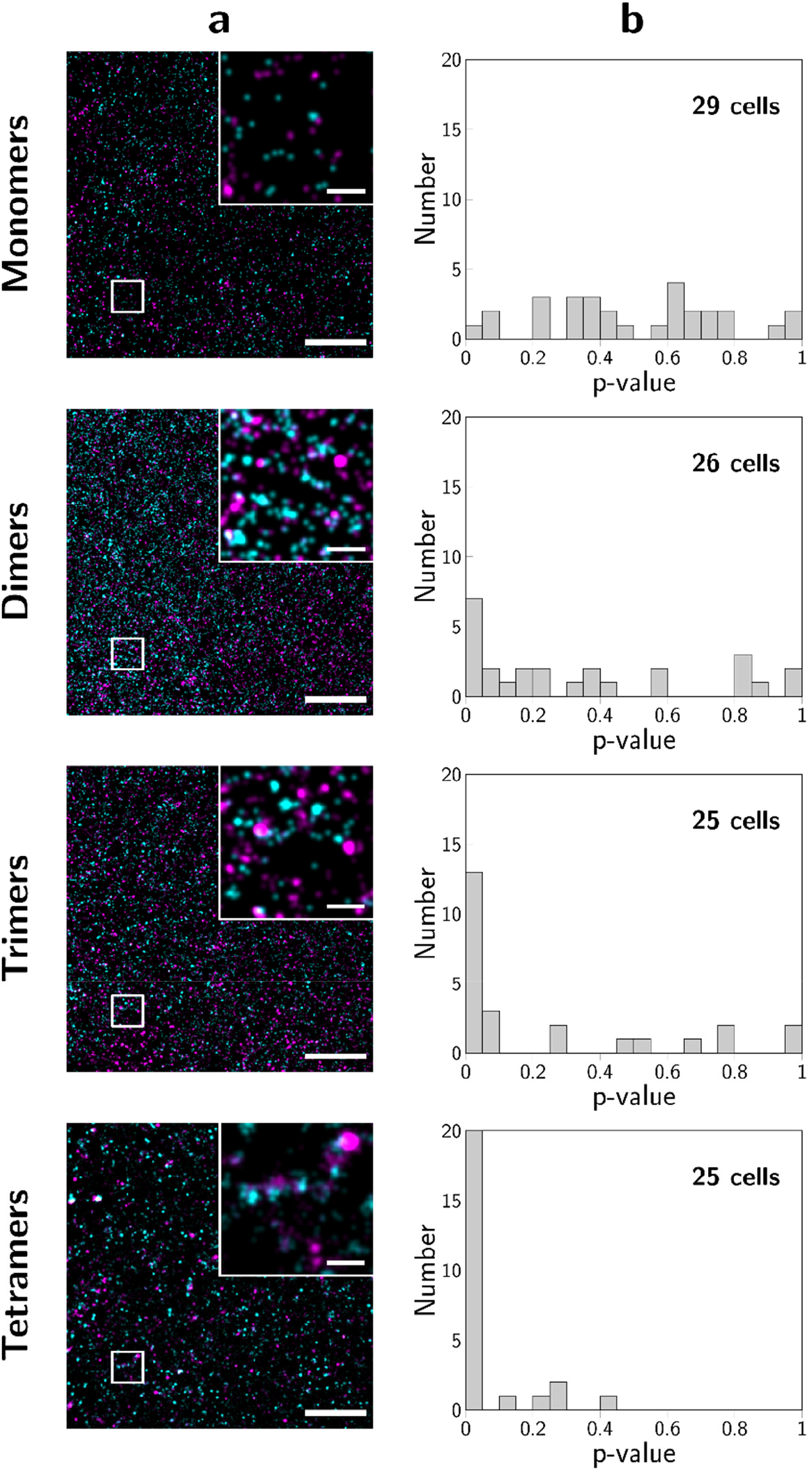
2-CLASTA analysis of an experimental data set. We analyzed GPI-anchored concatemers of SNAP-tags with n=1 to 4 subunits expressed in HeLa cells as mimicry of n-mers. For dSTORM experiments, cells were labeled with SNAP_488_ and SNAP_647_. Panels (**a**) show two-color localization maps for representative cells, and panels (**b**) histograms of p-values obtained from at least 4 independent experiments per n-mer. Scale bars 250 nm (inset) and 2 μm.

There is, however, a non-negligible fraction of cells which show p-values>0.05, even in the case of oligomeric SNAP constructs. This effect is rather prominent for dimers, and decreases with increasing degree of oligomerization. In a practical situation, however, one should note that different cells show different protein expression levels, thereby yielding a variability in the number of molecules within the region of interest. As shown in Fig. 2a, a low number of molecules would reduce the sensitivity for the detection of oligomers, or – in other words – would likely yield a high p-value. Indeed, when plotting the obtained p-value versus the number of localizations obtained per cell, we found a trend for high p-values at low localization numbers, which became more pronounced with increasing degree of oligomerization (**Fig. S10**). Particularly, for appr. 1,000 molecules per image – corresponding to appr. 5,000 localizations, we expect reduced sensitivity, which agrees with **Fig. S10**.

## Discussion

We present here a parameter-free method to statistically assess the question whether biomolecules are distributed randomly on a two-dimensional surface, yielding a p-value as output parameter. The method is compatible with most fluorescence labelling techniques, as long as it is ensured that each protein molecule is connected to one color channel only: this includes fluorescent antibodies or nanobodies, tags, or low affinity binders^16^.

Particularly, two challenges have to be approached in nanocluster analysis.

i. Obtaining the localization maps of a truly random protein distribution as a reliable standard for comparison with the experimental data. If such a distribution was available, comparative analysis such as Rényi divergence^17^ would be feasible. It turns out, however, that localization maps -as they would result from a truly random biomolecular distribution - are difficult to obtain, particularly since the photophysics of organic dyes often changes with the local environment of the chromophore^18^. We circumvented the problem by analyzing not the images themselves, but a correlation metric between the localizations of the two color channels (in our case, the nearest neighbor distances). In principle, also other metrics could be used for the significance test (e.g. pair cross-correlation analysis^7,19^ or Ripley’s covariate analysis^20^, especially for testing deviations on length scales beyond the nearest neighbors.
ii. Interpreting the results in statistical terms. The chosen analysis strategy based on correlation metrics offers the advantage that potential correlations between the two color channels can be deliberately broken, here by applying a toroidal shift to one of the two color channels. By this, the univariate spatial structure of each localization pattern is conserved, while any possible correlations are removed. This provides the possibility of significance testing between the original data and the randomized control data sets as an additional advantage.

To make the method immediately applicable, we provide a plugin for ImageJ (see **Supporting Material**). The experimental basis is a chromatically corrected two-color SMLM data-set analyzed by standard single molecule localization tools^21^.

In the following, we give a brief discussion on the strengths and potential pitfalls of our approach:

### Strengths

i. 2-CLASTA is stable against mistakes in chromatic or drift correction. As long as errors are smaller than typical cross-correlation distances of the two color-channels, the effects on the obtained p-values are marginal.
ii. 2-CLASTA is not impaired by blinking dye molecules, and does not require the recording of single molecule blinking statistics (as e.g. in the methods published in references^5,10,11^), making it insensitive to overcounting problems. In addition, 2-CLASTA can directly be applied to single images, thereby simplifying experimental efforts compared to our previously published method of label density variation^8^.
iii. The sensitivity of 2-CLASTA is not affected by any unknown characteristics of the clusters. No assumptions on cluster parameters (size, shape, occupancy) are required for the test. The test performs well even down to the detection of dimers, reflecting the smallest possible clusters.
iv. 2-CLASTA is stable against real live experimental challenges: A typical experiment contains non-specific localizations, or false negatives as a consequence of insufficient degree of labeling. Also the labeling ratio of the two colors may be unbalanced. We extensively tested the influence of such issues in Monte Carlo simulations, and found that the test is very robust over a wide range of parameters.

### Potential pitfalls

The sample topography may influence the obtained results: Without further information, it is reasonable to assume a completely random distribution of biomolecules on a flat two-dimensional surface parallel to the focal plane as the null hypothesis of the test. Randomly distributed biomolecules on an arbitrary two-dimensional manifold, however, may lead to virtual clustering in the projection onto a two-dimensional plane. For example, invaginations of the plasma membrane or cell borders will cause the accumulation of the detected positions of membrane proteins in the 2D projection^22^, and hence will likely lead to a rejection of the null hypothesis. In principle, such situations can be identified by analyzing the localization distribution in 3D.

## Conclusion

Taken together, we believe that the 2-CLASTA approach is well suited for a first assessment of spatial biomolecular distributions, before more sophisticated methods are used to characterize the clustering quantitatively^6^. By providing p-values, it makes use of the appropriate statistical parameter to test whether a specific data set is in agreement with a particular hypothesis^23^. Here, small p-values indicate suspicious deviations from randomness. Large p-values, in contrast, do not indicate any peculiarities in the sample; most notably, they do not prove a spatially random distribution of biomolecules. One should note that care has to be taken when interpreting the results of significance tests^23,24^. As a particular example, fishing for data sets with small p-values should be avoided.

A further application of 2-CLASTA is the analysis of co-localization of two different types of biomolecules: in this case, the two colors would be used to target the two different biomolecules. In this paper, we provide the framework to test for biomolecular association: extension towards assessment of biomolecular repulsion is straightforward and described in the Methods section.

## Materials and Methods

### Cell culture, DNA constructs, and reagents

All chemicals and cell culture supplies were from Sigma if not noted otherwise. All reagents for molecular cloning were from New England Biolabs. HeLa cells were purchased from DSMZ (ACC 57 Lot 23) and cultured in DMEM high glucose medium (D6439) supplemented with 10% fetal bovine serum (F7524) and 1 kU/ml penicillin-streptomycin (P4333). All cells were grown in a humidified atmosphere at 37°C and 5% CO_2_.

For transient transfection of HeLa cells with GPI-anchored SNAP concatemers, we fused one or multiple copies of the SNAP_f_ sequence to the N-terminus of the GPI-anchor signal of the human folate receptor. To this end, we carried out PCR to amplify the SNAP_N9183S_ sequence from pSNAP_f_ (N9183S) with >15 nt overhangs complementary to adjacent regions of the following SNAP_f_ copy. We then used the Gibson assembly Master Mix (E2611) following the supplier’s instructions to iteratively insert multiple consecutive copies of the SNAP_f_ sequence in frame with the GPI anchor. The resulting colonies were screened by site specific restriction digest using HindIII (R3104) to verify the number of inserted copies.

SNAP-Surface^®^ Alexa Fluor^®^ 488 (SNAP_488_) and SNAP-Surface^®^ Alexa Fluor^®^ 647 (SNAP_647_) were from New England BioLabs. Both labels were reconstituted in water-free DMSO (276855) at 10mg/ml, aliquoted and stored at −20°C until used.

STORM blinking buffer consisted of PBS, 50mM β-Mercaptoethylamine (30070), 3% (v/v) OxyFluor™ (Oxyrase Inc., Mansfield, Ohio, U.S.A.), and 20% (v/v) sodium DL-lactate (L1375)^25^. The pH was adjusted to 8-8.5 using 1M NaOH.

### Sample preparation

Cells were transfected by reverse transfection using Turbofect (ThermoFisher, R0531) according to the supplier’s instructions with Opti-MEM (Gibco, 31985062) as serum-free growth medium. Briefly, cells were detached from tissue culture flasks using Accutase (A6964). Subsequently, approximately 50,000 cells were mixed with Turbofect-DNA complexes and seeded on fibronectin-coated (F1141) LabTek chambers (Nunc) and incubated overnight. The following day, cells were labeled for 30-45 min in the incubator with 50nM SNAP_488_ and 1μM SNAP_647_ diluted in cell culture medium. After labeling, cells were extensively washed with HBSS, and fixed with 4% formaldehyde (Thermo Scientific, R28908) and 0.2% glutaraldehyde (GA) for 30 min at room temperature. After another series of two washing steps, we added 450μl freshly prepared STORM buffer immediately prior to imaging.

### Superresolution microscopy and image reconstruction

A Zeiss Axiovert 200 microscope equipped with a 100x Plan-Apochromat (NA=1.46) objective (Zeiss) was used for imaging samples in objective-based total internal reflection (TIR) configuration. TIR illumination was achieved by shifting the excitation beam parallel to the optical axis with a mirror mounted on a motorized table. The setup was further equipped with a 640 nm diode laser (Obis640, Coherent), a 405 nm diode laser (iBeam smart 405, Toptica) and a 488 nm diode laser (iBeam smart 488, Toptica). Laser lines were overlaid with an OBIS Galaxy beam combiner (Coherent). Laser intensity and timings were modulated using in-house developed LabVIEW software (National Instruments). To separate emission from excitation light, we used a dichroic mirror (Z488 647 RPC, Chroma). Images were split chromatically into two emission channels using an Optosplit2 (Cairn Research) with a dichroic mirror (DD640-FDi01-25×36, Semrock) and additional emission filters for each color channel (690/70H and FF01-550/88-25, Chroma). All data was recorded on a back-illuminated EM-CCD camera (Andor iXon DU897-DV).

Typically, we recorded sequences of 20 000 frames in alternating excitation mode. Samples were illuminated repeatedly at 640 nm, 405 nm, and 488 nm with 2-3 kW/cm^2^ intensity (640 nm and 488 nm) and 3-5 W/cm^2^ (405 nm); intensities were measured in epi-configuration. We selected the illumination times in ranges of 3 ms – 10 ms (640 nm), 3 ms – 30 ms (488 nm), and 6 ms (405 nm). Time delays between consecutive illuminations were below 6 ms. The camera was readout after the 640 nm and after the 488 nm illumination, yielding 10 000 frames in each color channel. Only data from those frames were included in the analysis, in which well-separated single molecule signals were observable.

We recorded calibration images of immobilized fluorescent beads after each experiment (TetraSpeck Fluorescent Microspheres, life technologies, T14792) and registered the images as described previously^26^. Single molecule localization and image reconstruction was performed using the open-source imageJ plugin ThunderSTORM^27^.

### Calculation of p-values

We compared the positions of all localizations obtained in the red color channel (*x_red_,y_red_*) with those obtained in the blue color channel (*x_blue_,y_irtue_*). For this, we calculated the distribution of distances, r, from each red localization to the nearest blue localization, and determined its cumulative distribution function *cdf*(*r*). To determine the distribution of nearest neighbor distances under the null model we applied a toroidal shift^14^ to the positions of the red color channel, according to 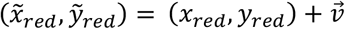, where 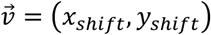 is the shift vector, with periodic boundary conditions set by the region of interest. The according nearest neighbor distribution was calculated as described, yielding *cdf_rand_*(*r*). The toroidal shift was repeated N-times with random shift vectors 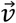 chosen uniformly within the region of interest, yielding *N* realizations of the null model of a random distribution of biomolecules. To compare the distributions, we first calculated 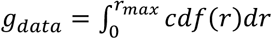 and the set 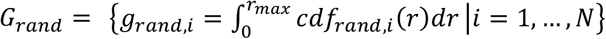. We next determined *rank*(*g_data_, G*), which is defined as the rank of *g_data_* within the set union *G = G_rand_* ∪ {*g_data_*}, where the statistical rank is measured in descending order. Finally, 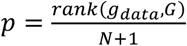 yields the one-sided p-value^28^. Naturally, this value is limited to discrete numbers with steps of 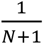. If not mentioned otherwise we chose *r_max_* → ∞. For practical reasons, we set in this case *r_max_* to the maximum nearest neighbor distance occurring during the whole analysis. In principle, the method can also be used to test for biomolecular repulsion; in this case, *g_data_* needs to be ranked within *G* in ascending order for calculation of the p-value.

### Simulations

Conceptually, simulations were performed as described previously^15^.

First, we simulated the underlying protein distributions for regions of 10 x 10 μm^2^, reflecting approximately the size of a typical cell. For all simulations we used 75 molecules per μm^2^, if not mentioned otherwise.

#### Simulation of oligomers

we distributed oligomers randomly within the region of interest, and assigned n biomolecules to each n-mer position (n=1 to 4). A random distribution of biomolecules is naturally reflected by the case of n=1.

#### Simulation of areas of enrichment or depletion of biomolecules

Circular domains with a radius of 20, 40, 60, 80, 100 or 150 nm were distributed randomly onto the region of interest with adjustable number of domains per μm^2^ (3, 5, 10, 15, 20 and 25). The number of biomolecules per domain was calculated from the total number of simulated molecules (here 7,500), the fraction of molecules inside domains (20, 40, 60, 80, 100 %), and the number of simulated domains, assuming a Poissonian distribution. Biomolecules were distributed randomly within the domains. The remaining molecules were distributed randomly in the areas outside of the domains.

Second, two different types of labels, corresponding to the two colors, were assigned randomly to the molecules according to the specified labeling ratio, assuming Binomial statistics.

Third, to simulate blinking, we assigned a number of detections to each label. This number was drawn from empirical probability distributions recorded at low labeling concentrations in dSTORM experiments on HeLa cells expressing GPI-anchored SNAP-tag monomers and labelled with SNAP_488_ or SNAP_647_ (**Fig. S1**) For the simulations shown in **Fig. S3 and Fig S9.**, we used blinking statistics determined previously^15^. Localization errors were simulated by spreading these detections using a Gaussian profile centered on the molecule position with a width of 30 nm, which corresponds to typical localization errors achieved in SMLM experiments. We assumed identical localization errors for the two color channels.

Fourth, to account for experimental errors in the “realistic” scenarios, we included unspecifically bound labels at a mean density of 5 labels/μm^2^ for each color channel, assuming the blinking statistics determined for SNAP_488_ and SNAP_647_. We finally considered also false positive localizations by adding a background of 1 (2) signals/μm^2^ for the red (blue) color channel, again with experimentally determined blinking statistics obtained in unlabeled cells.

Fifth, to account for stage drift in **Fig. 2d** we assumed alternating laser excitation and hence added a global drift vector 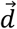 to the localizations of both color channels obtained at time *t* according to 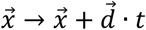.

If not mentioned otherwise, 100 simulations were performed for each experimental condition.

If not mentioned otherwise, we used the following set of parameters: 10 x 10 μm^2^ region of interest, 75 molecules per μm^2^, a balanced labeling ratio between the two color channels, no stage drift, 30 nm localization error (standard deviation). For the “ideal” scenario we simulated 100 % labeling efficiency, no unspecifically bound labels and no unspecific background signals. For the “realistic” scenario we simulated 40 % labeling efficiency, 5 unspecifically bound labels per μm^2^ and color channel, and 1 or 2 unspecific background signals per μm^2^ in the red and blue color channel, respectively.

## Acknowledgements

This work was supported by the Austrian Science Fund with project numbers F 6809-N36 and P 26337-B21 (to G.J.S.), P 27941-E28 (to F.B.), and a DOC Fellowship (24793) from the Austrian Academy of Sciences (A.M.A.). G.J.S and R.S. acknowledge support by the PhD program “MEIBio (Molecular and Elemental Imaging in Biosciences)” provided by TU Wien.

## Supporting Material

Supporting Information including 10 supplementary figures is available free of charge. We provide the code for data analysis with 2-CLASTA as ImageJ plugin. The Supplementary Software and associated manuals are available free of charge on https://github.com/schuetzgroup/2-CLASTA.git or https://owncloud.tuwien.ac.at/index.php/s/qwCyE22p5tk6IyY. All Matlab code is available from the corresponding authors upon reasonable request.

## Data availability

The data that support the findings of this study are available from the corresponding authors upon reasonable request.

## Author contributions

A.M.A. and M.C.S. contributed equally. F.B. and G.J.S. conceived the study; A.M.A., F.B., G.J.S. and M.C.S., designed the analytical method; M.C.S. developed the algorithm for p-value calculation; A.M.A. and M.C.S. wrote the code for the analytical methods and the simulations; A.M.A., F.B., M.B. and M.C.S. performed experiments; A.M.A., M.B., and M.C.S. analyzed the data; C.H. wrote the ImageJ plugin; G.J.S. and R.S. directed research; A.M.A., G.J.S. and M.C.S. wrote the manuscript.

