## Supplementary Figures S1 - S10 for "Verifying molecular clusters by 2-color localization microscopy and significance testing"

### Running title: Verifying molecular clusters by 2-CLASTA

**Keywords**: Superresolution microscopy, significance test, nanocluster, fluorophore blinking, single molecule localization microscopy, protein oligomers, two-color STORM

[1] Both authors contributed equally

[2] Institute of Applied Physics, TU Wien, Getreidemarkt 9, A-1060 Vienna, Austria.

[3] Institute of Visual Computing and Human-Centered Technology, TU Wien, Favoritenstrasse 9-11, A-1040 Vienna, Austria

[*] Corresponding author

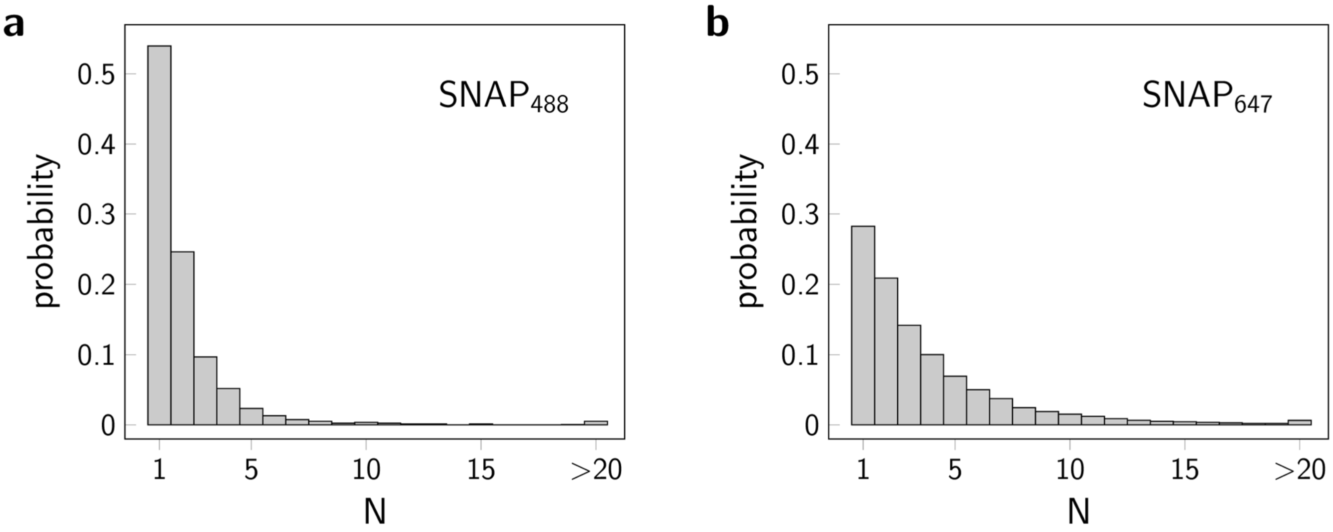


**Figure S1**

**Single molecule blinking statistics for SNAP-labels**

Blinking statistics of individual $SNAP_{488}$ (**a**) and $SNAP_{647}$ molecules (**b**) were recorded on fixed HeLa cells expressing the monomeric SNAP-GPI protein construct. Cells were labelled at sufficiently low concentrations of the SNAP-ligand so that well-separated single molecule signals could be observed. Histograms show the probability of a single label to result in N localizations.

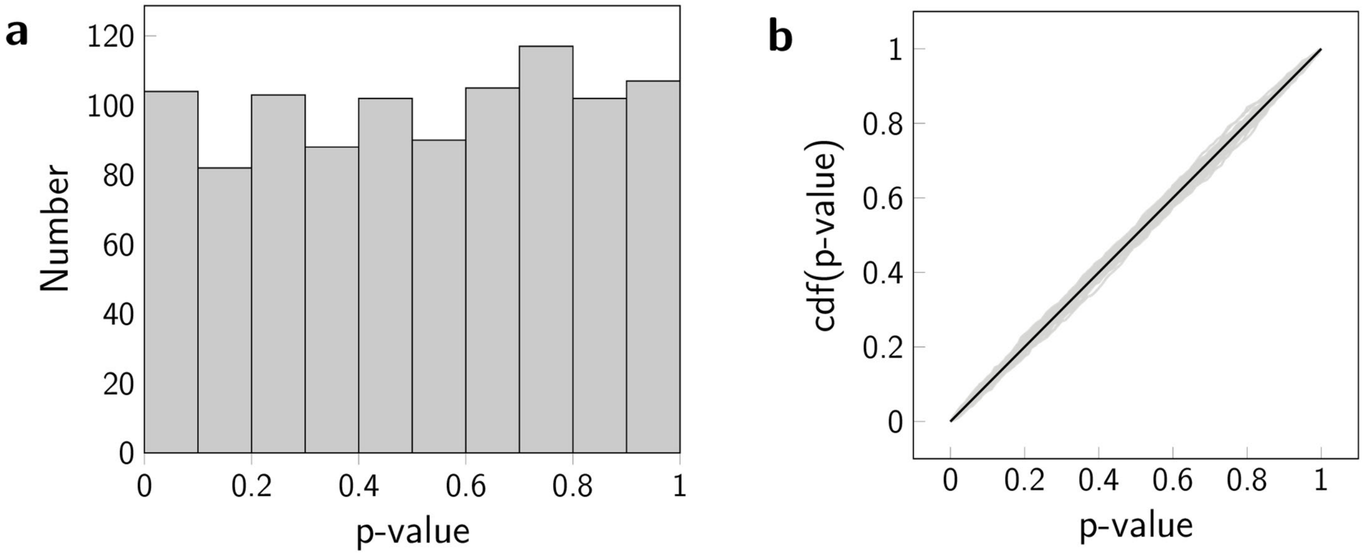


**Figure S2**

**Testing realizations of the null hypothesis**

1000 realizations of the null hypothesis of randomly distributed biomolecules were simulated and tested with 2-CLASTA. We simulated 36 different parameter sets, varying the number of molecules (50; 75; 100 molecules per µm²), the labeling efficiency (40%, 60%, 80% and 100%) and the labeling ratio (3:7, 2:3, 1:1). All other simulation parameters were held constant. (**a**) Histogram of the p-values obtained for one exemplary parameter set. (**b**) Cumulative distribution functions of p-values for various parameter settings are shown in light gray. The ideal uniform distribution is indicated by the solid black line, as comparison.

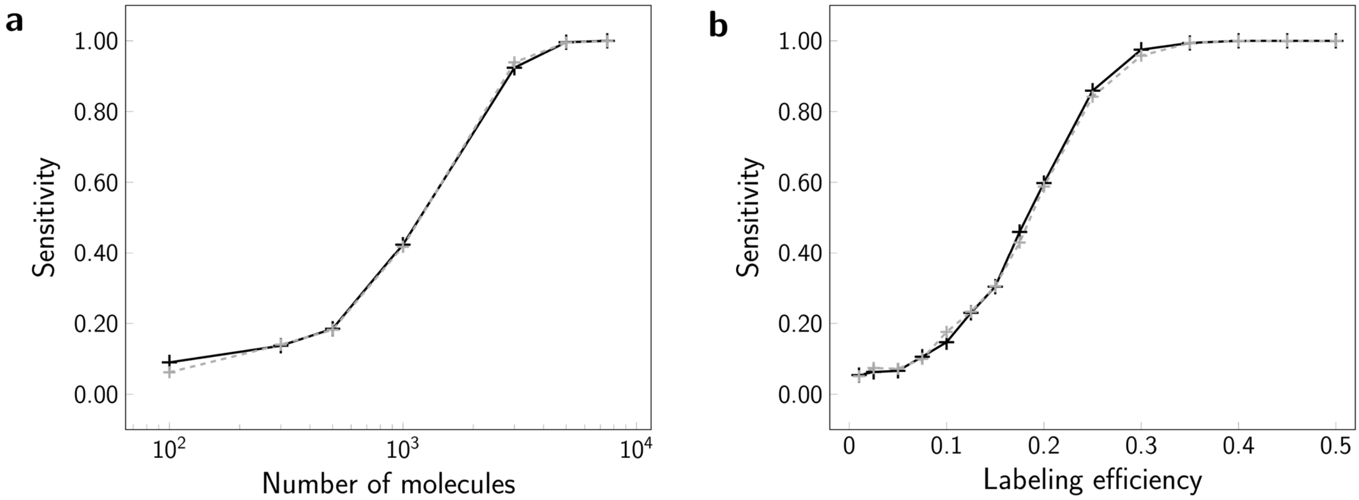


**Figure S3**

**Influence of different fluorescent labels on 2-CLASTA sensitivity**

We determined the sensitivity as a function of the number of molecules (**a**), and the labeling efficiency (**b**), for the case of molecular dimers, assuming the “realistic” scenario (**+**). Included are experimentally derived blinking statistics for $SNAP_{488}$ and $SNAP_{647}$ (black), and for $KT3_{647}$ and PS-CFP2 (gray). If not varied in the respective subpanel, parameters in all simulations were set to a molecular density of 75 molecules/µm², a labeling efficiency of 40%, a labeling ratio of 1:1, and no stage drift.

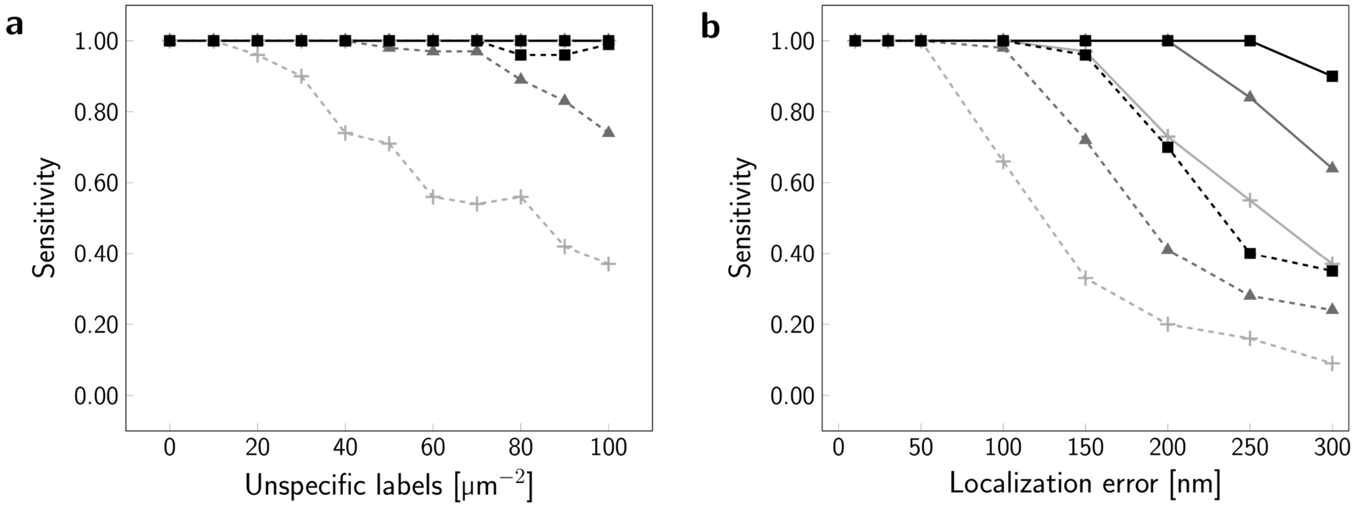


**Figure S4**

**Influence of unspecifically bound labels and localization errors on the sensitivity**

The sensitivity of 2-CLASTA was determined as a function of unspecifically bound labels (**a**), and for different localization errors (**b**). We simulated dimers (**+**), trimers (▲) and tetramers (■), both for the “ideal” (solid line) and the “realistic” scenario (dashed line). Each data point corresponds to 100 independent simulations.

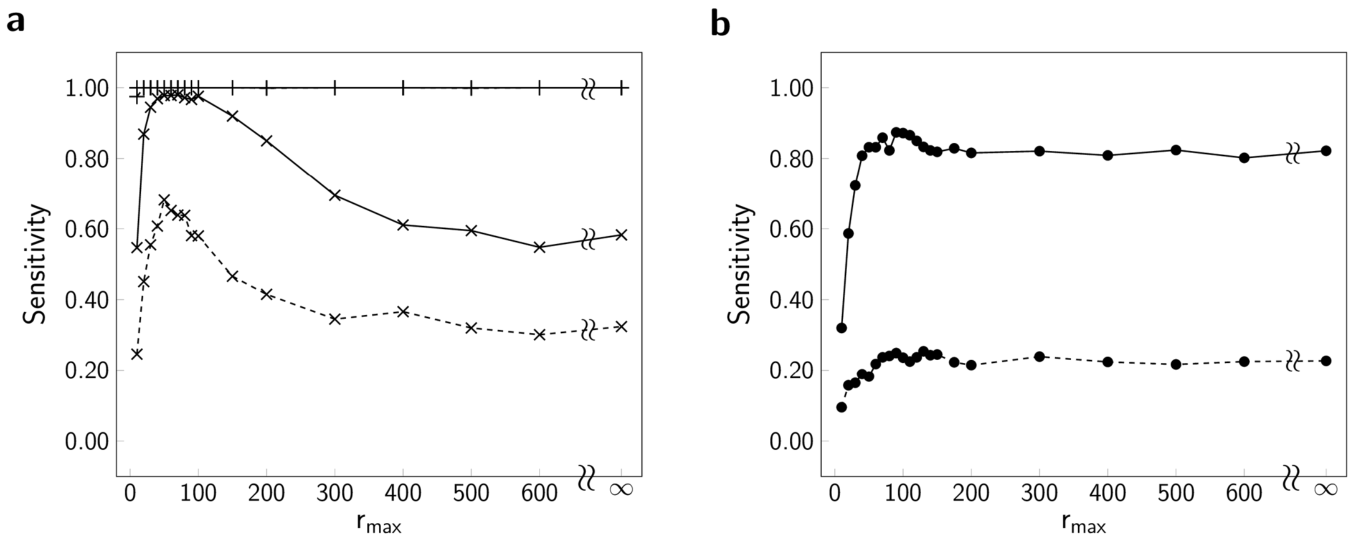


**Figure S5**

**Influence of $r_{max}$ on 2-CLASTA sensitivity**

(**a**) Influence of the analysis parameter $r_{max}$ on 2-CLASTA sensitivity for the detection of biomolecular dimers, both for the “ideal” (solid line, **+**) and the “realistic” scenario (dashed line, **+**). In addition, we also included simulations for cases in which we reduced the labeling efficiency to 15%, while keeping all other parameters as before (x). (**b**) Influence of the analysis parameter on 2-CLASTA sensitivity for the detection of circular nanodomains of 100 nm radius, 3 clusters per µm² and 20% of molecules inside the nanodomains, both for the “ideal” (solid line, ●) and the “realistic” scenario (dashed line, ●). Each data point corresponds to 1000 independent simulations.

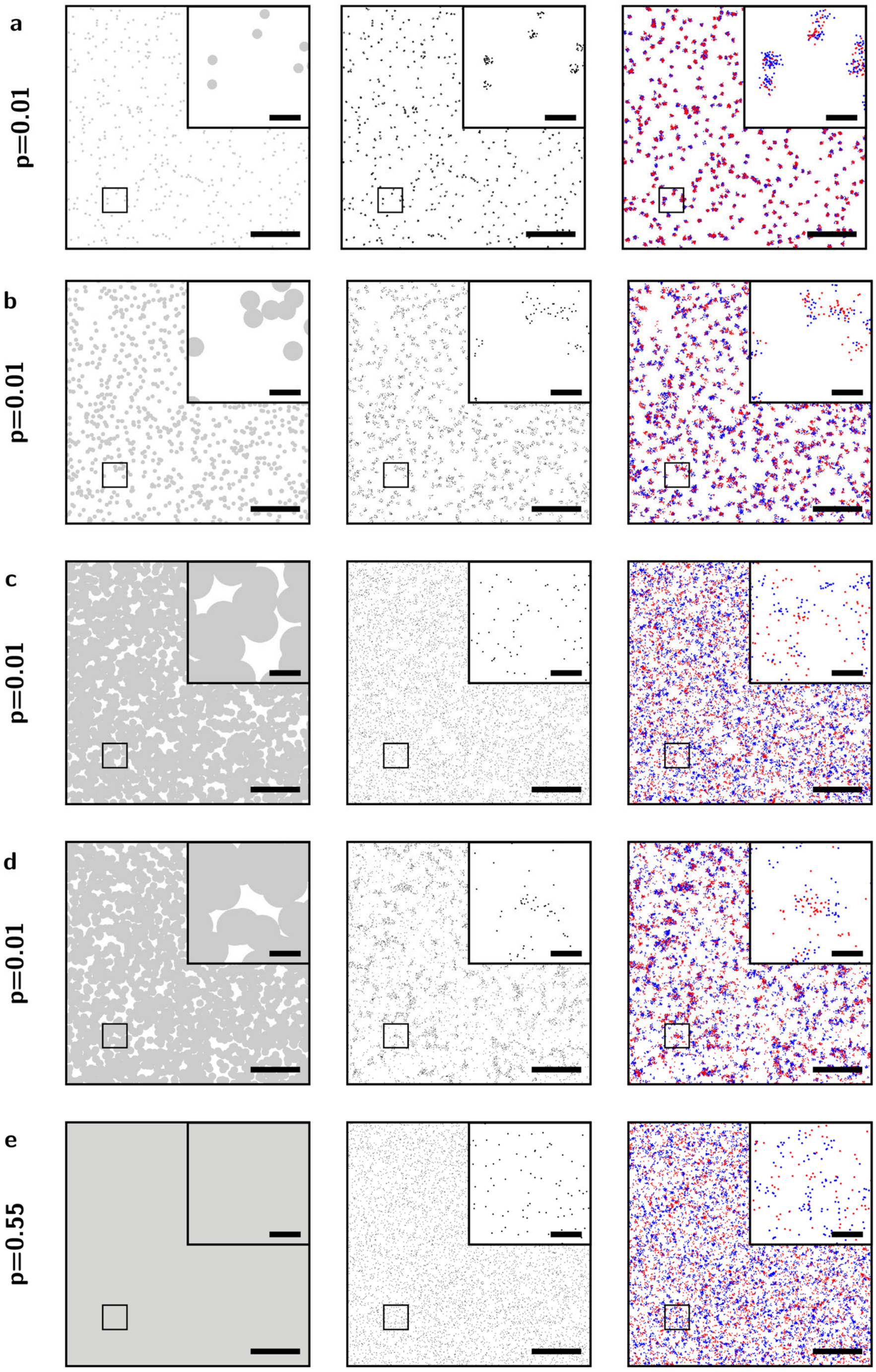


**Figure S6**

**Examples of simulated scenarios for nanodomains of biomolecules ("ideal" scenario)**

The underlying simulated nanodomains (left), the positions of simulated biomolecules (center), and the resulting localization maps (right) are shown for rare small clusters (**a**), medium clusters (**b**), large frequent clusters (**c**), exclusion areas (**d**), and a random distribution of biomolecules (**e**). The resulting p-value for each scenario is indicated on the left. Scale bars 250 nm (inset) and 2 µm.

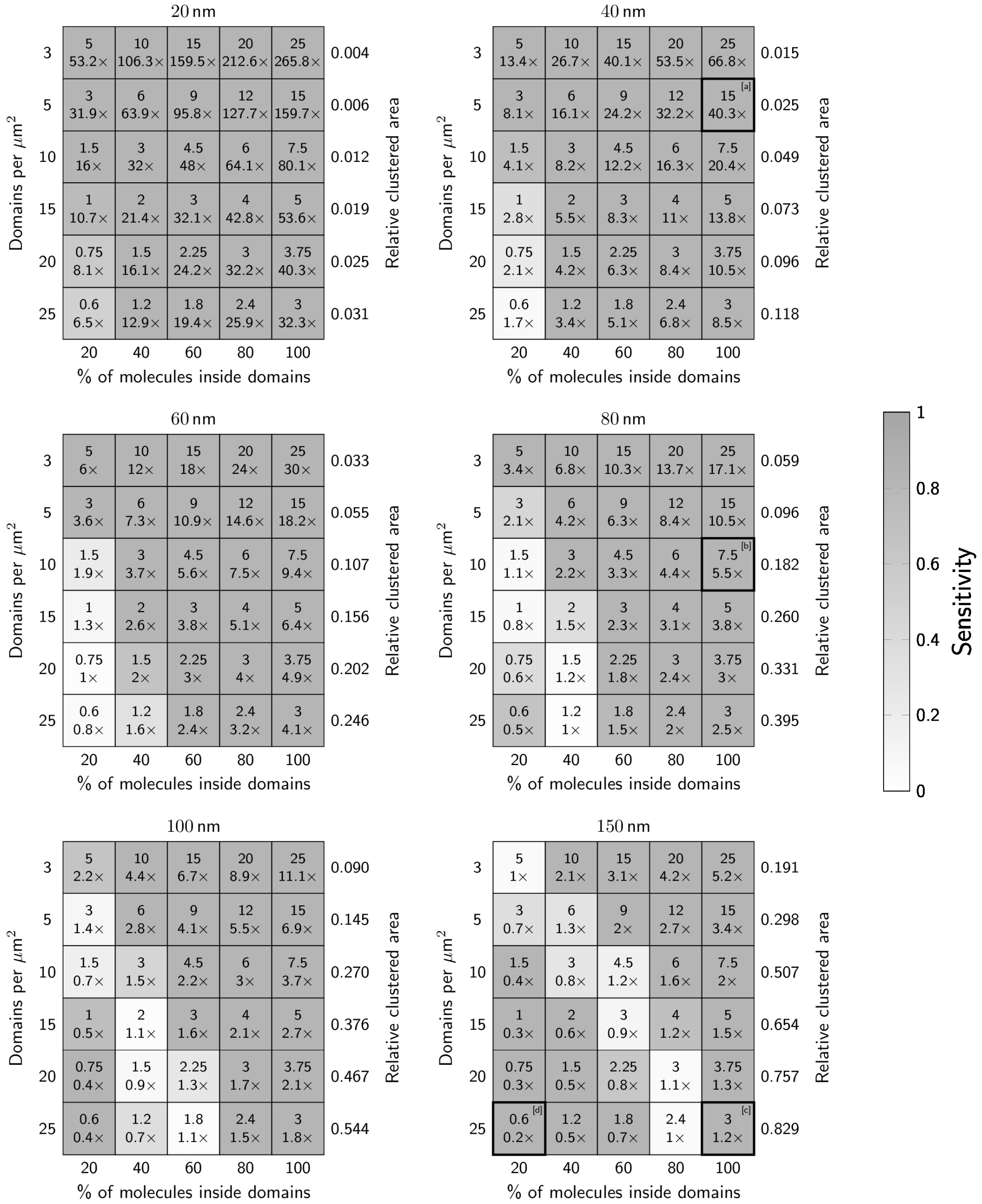


**Figure S7**

**Sensitivity of 2-CLASTA to detect protein enrichment or depletion ("ideal" case)**

Simulations were performed for circular nanodomains with radii of 20, 40, 60, 80, 100 and 150 nm. The number of domains per µm² was varied between 3 and 25, and the percentage of molecules inside the domains between 20% and 100%. Numbers in individual fields indicate the average number of molecules per domain, and the relative enrichment or depletion of molecules compared to a random distribution with identical average density. The gray scale indicates the fraction of scenarios with a p-value below the significance level $\alpha$=0.05, reflecting the sensitivity. Each field corresponds to 100 independent simulations. The bold black boxes indicate the scenarios shown in the exemplary images in **Fig. S6**, letters indicate subpanels.

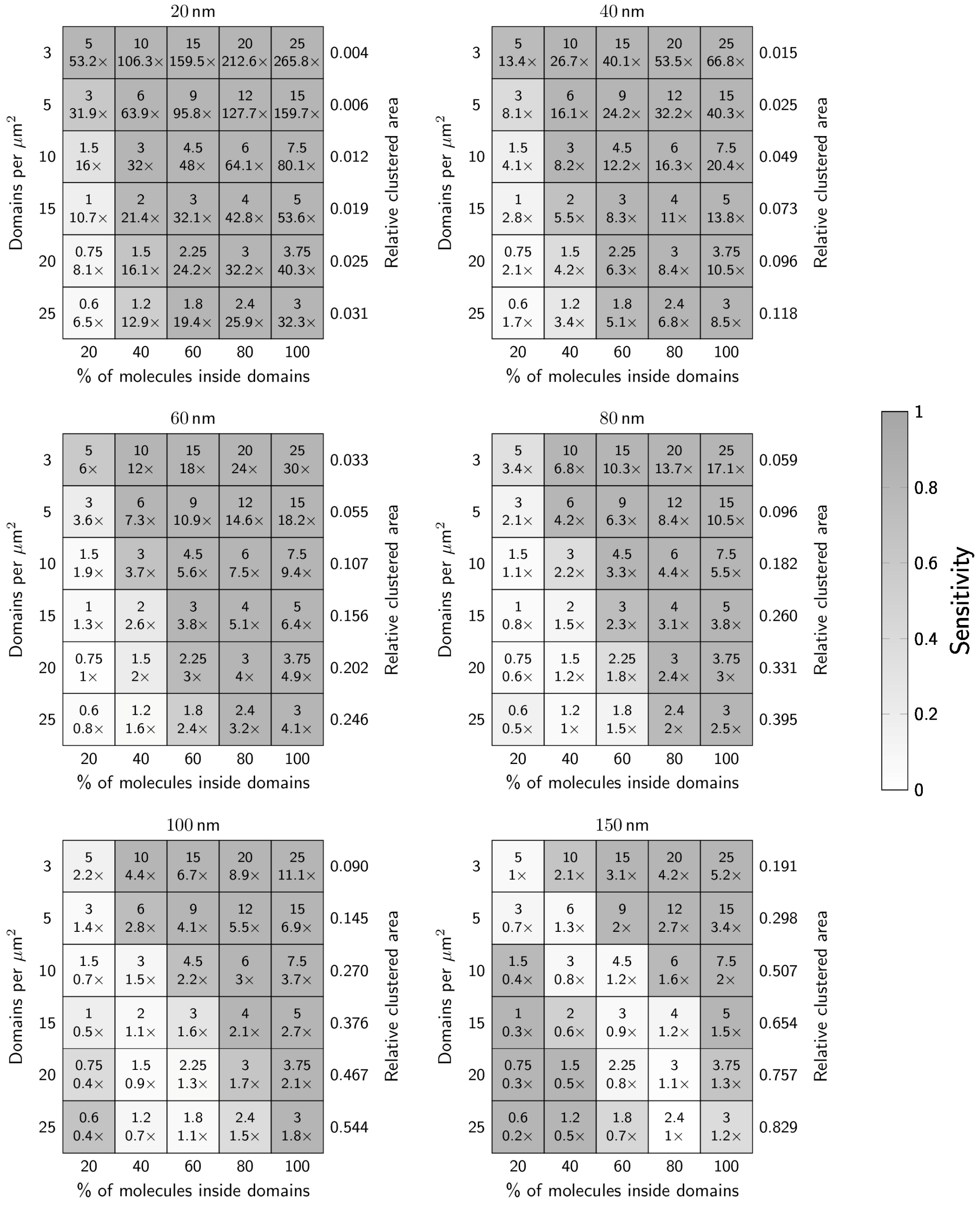


**Figure S8**

**Sensitivity of 2-CLASTA to detect protein enrichment or depletion (“realistic” case)**

Simulations were performed as described in Supplementary Figure 4. Number in individual fields indicate the average number of molecules per domain, and the relative enrichment or depletion of molecules compared to a random distribution with identical average density. The gray scale indicates the fraction of scenarios with a p-value below the significance level $\alpha$=0.05, reflecting the sensitivity. Each field corresponds to 100 independent simulations.

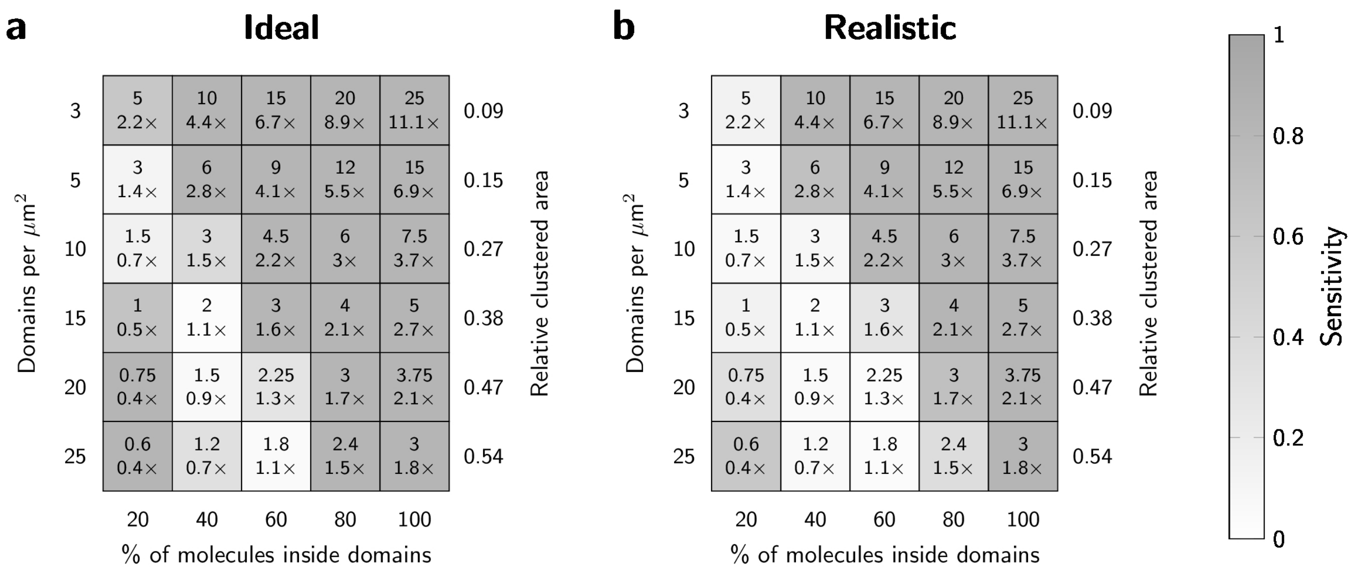


**Figure S9**

**Sensitivity of 2-CLASTA to detect protein enrichment or depletion using a different fluorescent label**

We determined the sensitivity of 2-CLASTA for varying densities of circular domains and percentage of molecules inside the domains, assuming the blinking statistics for $KT3_{647}$ and PS-CFP2 (gray). Data are shown for a cluster radius of 100 nm for the “ideal” case (**a**) and the “realistic” case (**b**). Number in individual fields indicate the average number of molecules per domain, and the relative enrichment or depletion of molecules compared to a random distribution with identical average density. The gray sale indicates the fraction of scenarios with a p-value below the significance level $\alpha$=0.05, reflecting the sensitivity. Each field corresponds to 100 independent simulations.

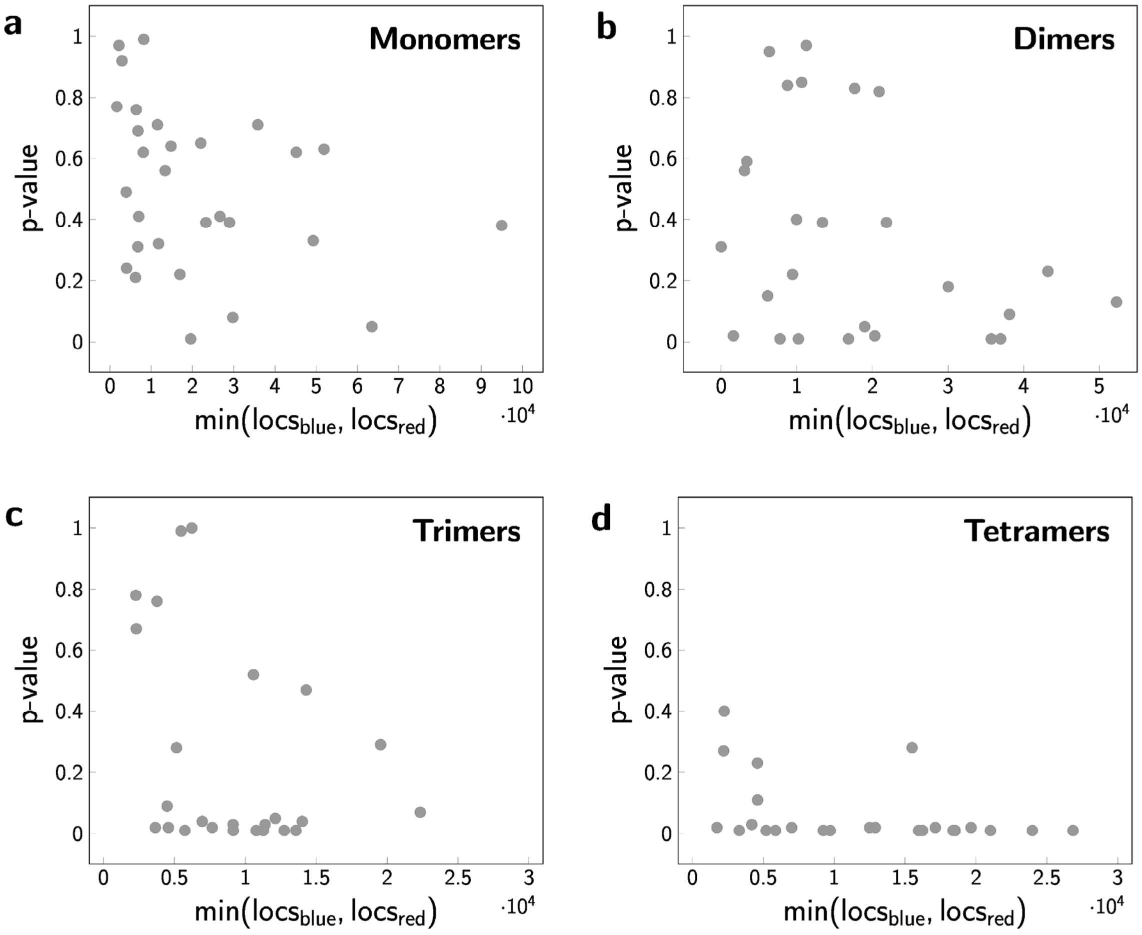


**Figure S10**

**Correlation of the obtained p-values versus the number of localization per region of interest**

We used the data sets shown in Fig.4. For each region of interest, we plotted the obtained p-value versus the minimum of the number of localizations recorded in the two color channels. Panels **a** - **d** correspond to monomers, dimers, trimers, and tetramers, respectively.
